# Towards understanding Afghanistan pea symbiotic phenotype through the molecular modeling of the interaction between LykX-Sym10 receptor heterodimer and Nod factors

**DOI:** 10.1101/2020.09.21.306514

**Authors:** Ya.V. Solovev, A.A. Igolkina, P.O. Kuliaev, A.S. Sulima, V.A. Zhukov, Yu.B. Porozov, E.A. Pidko, E.E. Andronov

**Affiliations:** Shemyakin-Ovchinnikov Institute of Bioorganic Chemistry, Russian Academy of Sciences, 117997, Moscow, Russian Federation; TheoMAT research group, ITMO University, Lomonosova St. 9, 191002 St. Petersburg, Russia; All-Russia Research Institute for Agricultural Microbiology (ARRIAM), Podbelsky Chaussee 3, Pushkin, 196608, Saint-Petersburg, Russian Federation; Saint-Petersburg State University, Saint-Petersburg, Russia; Sirius University of Science and Technology, Olympic ave. 1, 354340, Sochi, Russian Federation; I.M. Sechenov First Moscow State Medical University, Trubetskaya str. 8, Moscow 119991, Russian Federation; The laboratory of Bioinformatics, Department of Food Biotechnology and Engineering, ITMO University, Kronverksky pr. 49, lit. A, Saint Petersburg 197101, Russia; Delft University of Technology, Faculty of Applied Sciences, Inorganic Systems Engineering group, Department of Chemical Engineering, Van der Maasweg 9, 2629 HZ, Delft, The Netherland; V.V. Dokuchaev Soil Institute, Moscow, Russia

**Author notes:** Co-first authorship: Ya.V. Solovev and A.A. Igolkina.

**Keywords:** pea, plant–rhizobia symbiosis, LykX, Sym2, Nod factor, molecular dynamics, COSMO-RS model

## Abstract

The difference in symbiotic specificity between peas of Afghanistan and European phenotypes was interrogated using molecular modeling. Considering segregating amino acid polymorphism, we examined interactions of pea LykX-Sym10 receptor heterodimers with four forms of Nod factor (NF) that varied in natural decorations (acetylation and length of the glucosamine chain). First, we showed the stability of the LykX-Sym10 dimer during molecular dynamics (MD) in solvent and in the presence of a membrane. Then, four NFs were separately docked to one European and two Afghanistan dimers, and the results of these interactions were in line with corresponding pea symbiotic phenotypes. The European variant of the LykX-Sym10 dimer effectively interacts with both acetylated and non-acetylated forms of NF, while the Afghanistan variants successfully interact with the acetylated form only. We additionally demonstrated that the length of the NF glucosamine chain contributes to controlling the effectiveness of the symbiotic interaction. The obtained results support a recent hypothesis that the LykX gene is a suitable candidate for the unidentified Sym2 allele, the determinant of pea specificity towards *Rhizobium leguminosarum bv. viciae* strains producing NFs with or without an acetylation decoration. The developed modeling methodology demonstrated its power in multiple searches for genetic determinants, when experimental detection of such determinants has proven extremely difficult.

## INTRODUCTION

The symbiosis between leguminous plants (*Fabaceae*) and nodule bacteria (collectively called *rhizobia*) demonstrates an extremely high specificity. When the host plant interacts with a large number of soil microorganisms looking for an appropriate partner, it should be selective enough to avoid penetration of any pathogens, discriminate between different types of *rhizobia*, and allow symbiotic interaction with the most effective partner. Interactions between partners are carried out by a complicated interplay of signaling; however, several essential details of these molecular mechanisms are still under question.

The molecular crosstalk between partners starts at the early stages of the plant–rhizobia interaction and results in root nodule formation (Figure 1a, photos). Bacteria secrete molecules called Nodulation factors (Nod factors, NFs), which serve as the primary recognition “key” signal for a plant “lock”. The structure of NFs was discovered more than 30 years ago; this molecule generally consists of 3, 4 or 5 N-acelylglucosamines linearly oligomerized through (1,4)-β-linkages and decorated with a fatty acid residue and various small substitutions that play a crucial role in partner recognition (Denarie, 1996; Ritsema et al., 1996). All these decorations are genetically controlled and determine NF shapes, which are distinctive for different *rhizobia* species (or strains). Each rhizobial strain produces several slightly different NF molecules that may allow the strain to extend the spectrum of its potential hosts (Mergaert et al., 1997). In contrast, the beneficial strategy for plants is to narrow their symbiotic specificity and interact with reliable symbiotic partners only. The co-evolution of these strategies results in cross-inoculation groups (CIGs) (Baldwin et al., 1927; Sears and Carroll, 1927). This concept describes distinct groups, each containing both legumes and *rhizobia* strains, so that all plant–bacteria combinations within a group form effective and highly specific symbiotic relationships. Originally, CIGs were thought to be non-overlapped; each legume species and *rhizobia* strain was considered to belong to only one group (Baldwin et al., 1927; Sears and Carroll, 1927). However, further studies demonstrated that real boundaries between CIGs are blurred, and many cases of non-canonical interactions between legumes and *rhizobia* from different groups exist (Provorov, 1994; Viprey et al., 2000). Even within a particular CIG, one can find variations in the symbiotic efficiency (Provorov, 1994).

**Figure 1.**
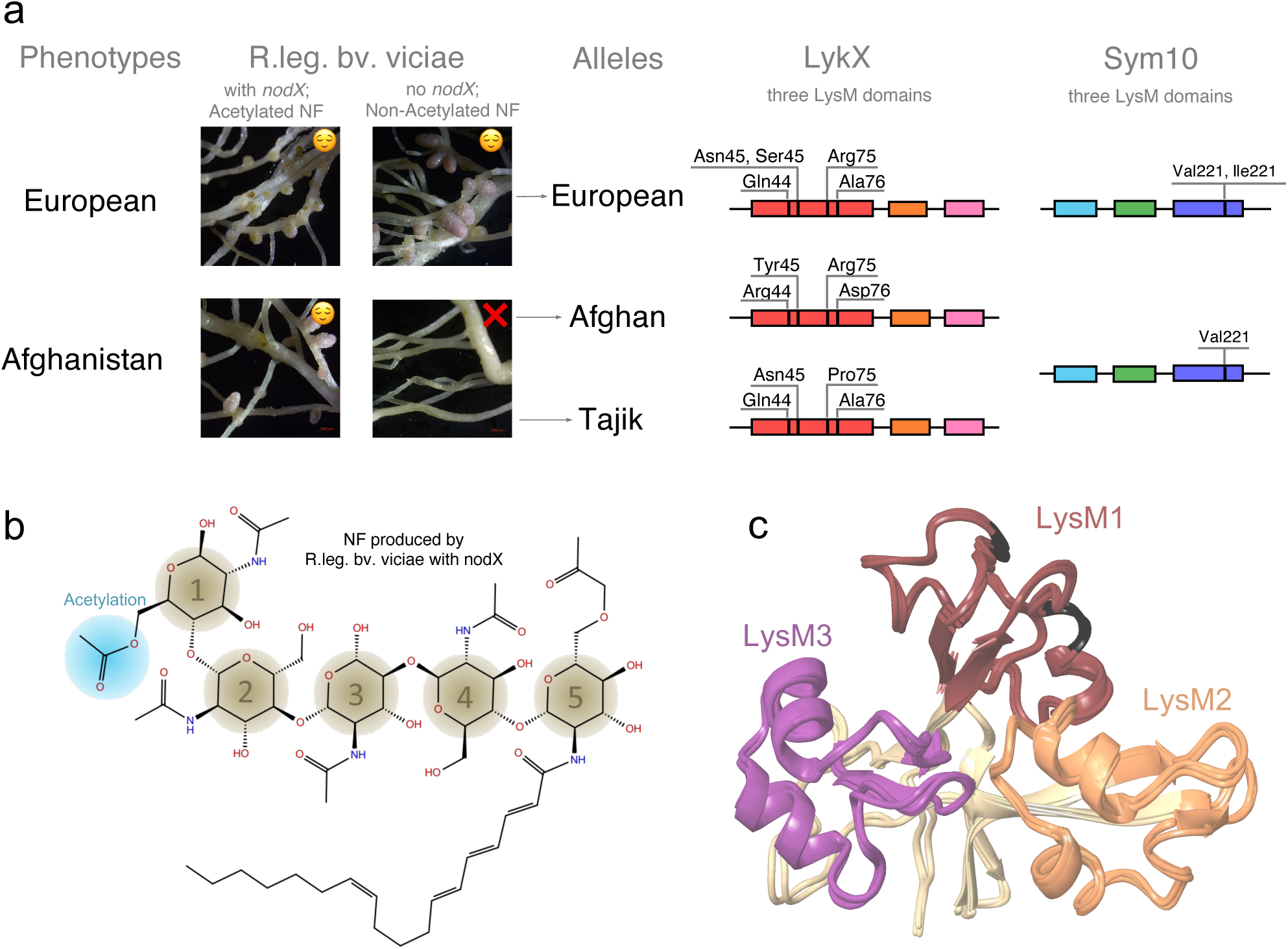
Phenotypes and alleles of garden pea (Pisum sativum L.). Garden pea has two symbiotic phenotypes (European and Afghanistan) differing in symbiosis with R.leg.bv.vicia, which have no *nodX* gene and produce non-acetylated NFs. At the molecular level, two alleles of LykX gene were found for Afghanistan phenotype (Afghan and Tajik). Three LysM domains and amino acid polymorphisms for LykX and Sym10 are shown. (b) Structure of a NF. (c) Structural alignment of the Afghan, Tajik, and European LykX proteins. LysM1 domains are colored in dark red (variable amino acids are highlighted with black), LysM2 – in faded orange, and LysM3 – in dark purple.

This variation within a CIG was demonstrated for the garden pea (*Pisum sativum L*.); Afghanistan and European pea landraces show different symbiotic success with *Rhizobium leguminosarum bv. viciae* depending on presence of the *nodX* gene in *rhizobia* (Firmin et al., 1993; Ovtsyna et al., 2007). This gene encodes an acetyltransferase that attaches a small acetylation decoration to NF (Figure 1b) (Davis et al., 1988; LIE, 1978; Ovtsyna et al., 1999). The Afghanistan pea landraces can interact only with *Rhizobium leguminosarum bv. viciae e* strains, which have the *nod*X gene, while European pea lines display relaxed specificity and can also be nodulated by strains lacking this gene (Figure 1a). The reason for this variation relates to the difference in the NF receptors of pea subtypes. However, the particular pea receptors that interact with NFs are not known.

Plant receptors for NF perception belong to a family of LysM domain-containing receptor-like kinases (LysM-RLKs), and contain three LysM domains in the extracellular part of the protein, a transmembrane domain, and an intracellular kinase domain (Bensmihen et al., 2011; Buendia et al., 2018). All plant LysM-RLKs fall into two classes: LYK (with active kinase) and LYR (with non-active pseudo-kinase) (Arrighi, 2006). A heterodimeric receptor of two LysM-RLKs from different classes is required for NF perception (Arrighi, 2006; Radutoiu et al., 2003; Zipfel and Oldroyd, 2017). The differences in receptor protein sequences, in theory, should reflect the difference in NF structure. However, in practice, different leguminous species with a relatively high level of between-species amino acid polymorphism in receptor genes effectively interact with the same rhizobial strains. Interestingly, a single amino acid variation in a particular LysM domain of the LysM-RLK NFR5 has been shown to dramatically change the recognition of NFs by Lotus species (Radutoiu et al., 2007). The latter example resembles the case of Afghanistan and European pea cultivars demonstrating a relatively low level of genetic divergence and, at the same time, clearly different symbiotic phenotypes (Figure 1a). To understand the molecular mechanisms of these different interactions, one needs to study NF–receptor interaction and the meaning of amino acid substitutions in this process. It is first necessary to interrogate what is known about the interactions between pea receptors and NF. Expectedly, NF receptors should be homologues of the previously identified legume LysM-RLKs, but in practice, it is almost impossible to match known pea LysM-RLKs with their homologues in other leguminous plants, as LysM-RLKs families consist of a number of closely related receptors.

The existence of the specific pea receptor for acetylated NF was shown with a classic genetic approach more than 20 years ago. It was well-proven that the difference in interaction exhibited by acetylated and non-acetylated NF is controlled by a single plant gene called Sym2 (Kozik et al., 1996, 1995); however, the position of this gene in the pea genome is unknown. Recently, the Sym2 candidate gene LykX (from LRY class) was proposed based on sequence comparison of receptor genes in Afghanistan and European pea subtypes (Sulima et al., 2017). Since NF perception requires a heterodimeric receptor, we considered the conventional Sym10 gene encoding its LYK subunit, and the LykX gene encoding an LYR subunit.

The most obvious way to prove the action of LykX as the NF receptor is by investigating the direct binding of the isolated receptor with NF. However, the evidence for direct binding of acetylated and non-acetylated NFs to LykX protein is still lacking. Only several attempts to show such binding have been successful to date (Broghammer et al., 2012; Kirienko et al., 2018). The difficulty of this detection may be caused by the fact that legume NF receptors work in heterodimers (Igolkina et al., 2018; Kirienko et al., 2019; Xiao et al., 2014), whereas modern technologies are aimed to test the “one receptor–one ligand” model. Additionally, NF isolation and purification are very time- and resource-consuming procedures, while the chemical synthesis of NF is extremely difficult due to the high complexity of its structure.

Therefore, as long as the demonstration of the physical interaction between receptor heterodimers and NFs remains challenging, molecular modeling is a suitable and appealing alternative to direct experimental approaches. Recently, based on this method, probable complexes of plant heterodimeric receptors with NF have been proposed, and the overlap between population polymorphisms in the contact zone between receptors in the complex has been analyzed (Igolkina et al., 2018). To date, molecular modeling provides the only opportunity for studying the effect of structural variations in NF and its receptors on the efficiency of their interactions.

In this study, we exploited the merits of molecular modeling to analyze the effect of specific chemical modifications in NF produced by *Rhizobium leguminosarum bv. viciae* on its interaction with the LykX-Sym10 receptor. For all possible LykX-Sym10-NF complexes, we performed two independent *in silico* validations — canonical MD and an original thermochemical-based pipeline — and obtained stable configurations in line with the observed pea phenotypes. The modeling analysis supported the LykX gene as a suitable determinant for Sym2.

## Methods

### Sequence analysis

We analyzed 95 amino acid sequences of *Ps*LykX receptor (MF155382–MF155469; MN187362– MN187364, MN200353–MN200358) (Sulima et al., 2019, 2017) and 8 amino acid sequences of *P*sSym10 receptor (MN727808, MN727809, MN727810, MN727811) (Sulima et al., 2019).

Among the LykX sequences used in this study, 7 represented the Afghanistan pea phenotype and 85 represented the European pea phenotype. According to the Sulima and colleagues, the Afghanistan pea phenotype has two alleles, Afghan and Tajik (Sulima et al., 2019), which were present in our dataset in 5 and 2 variants, respectively. Therefore, we separately analyzed three subsets of PsLykX sequences based on the following pea alleles: Afghan, Tajik, and European. It should be noted that the names of alleles do not reflect the geographical origin of the corresponding pea samples. Within 8 sequences of the PsSym10 receptor gene, 3 belonged to Afghanistan phenotype and 5 to the European phenotype (Supplementary Table 1).

Most of the analyzed sequences were obtained from previous studies, but four alleles of PsSym10 (MN727808, MN727809, MN727810, MN727811) were sequenced in the current study. DNA was extracted from young leaves (top or second-from-top node). Extraction was conducted according to the previously described CTAB protocol (Rogers and Bendich, 1985; Sulima et al., 2017). PCR was performed in 0.5 ml eppendorf-type microcentrifuge tubes on an iCycler (Bio-Rad, Hercules, CA, USA) or Dyad (Bio-Rad) thermocycler using the ScreenMix-HS kit (Evrogen, Moscow, Russia). The PCR cycling conditions were as follows: 95°C (5 min), 35 cycles [95°C (30 s), Tm (varying depending on primers) (30 s), 72°C (1 min)], 72°C (5 min). The PCR fragments were sequenced using the ABI Prism 3500xL system (Applied Biosystems, Palo Alto, CA, USA) at the Genomic Technologies, Proteomics, and Cell Biology Core Center of the ARRIAM (St. Petersburg, Russia). PCR primer sequences are listed in Supplementary Table 2. The resulting sequences have been deposited into the NCBI database (see accession in Supplementary Table 1).

Extra-membrane domains of LykX and Sym10 receptors contain 212 and 226 amino acids, respectively. Sequences for LykX and Sym10 were aligned separately in Mega X using the MUSCLE algorithm (Kumar et al., 2018). In each alignment, we analyzed polymorphic positions within and between allele subsets and focused only on polymorphic positions found between subsets. We hypothesized that the polymorphisms found between subsets are responsible for the difference in pea phenotypes.

### Protein homology modeling

Three LysM-containing kinase crystals were used as templates for the homology modeling of extracellular domains of the receptors: 5JCD of the *O. sativa* OsCEBiP chitin receptor (Liu et al., 2016), 4EBZ of the *A. thaliana* AtCERK1 chitin elicitor receptor kinase (Liu et al., 2012), and 5LS2 of the *L. japonicus* LysM-containing protein (Bozsoki et al., 2017).

For European and Afghan alleles of the LykX and Sym10 proteins, the extracellular and transmembrane domains were predicted with three separate methods: SWISS-MODEL (Waterhouse et al., 2018), Iterative Threading ASSEmbly Refinement (I-Tasser) (Zhang, 2009), and Phyre2 (Kelley et al., 2015) algorithms. The structure of the Tajik allele of LykX was obtained by a single amino acid mutation (ARG75PRO) in the European model and then relaxed using the MM approach. For all 5 obtained models (3 for LykX and 2 for Sym10), we analyzed Ramachandran plots and RMSD deviations from the corresponding template geometry, and filtered out models with defective secondary structures of the LysM domains. Final models were prepared using Protein Preparation Wizard (Madhavi Sastry et al., 2013; Schrödinger, 2018) (Sastry *et al*., 2013; Schrödinger, 2018) and minimized in Maestro using OPLS3ext force field (Roos et al., 2019).

### Protein–protein docking and clustering of dimers

For LykX-Sym10 pairs, we performed the protein–protein docking assay in the Piper package (Kozakov et al., 2006) and obtained 30 docking poses of dimers for each of the three pea alleles (European, Afghan, and Tajik).

Then, we clustered 90 obtained dimers (30×3) in the following way: using the Procrustes analysis (rotation and shift, without scaling) (Rudemo, 2000), we placed all dimers into the comparable coordinates, so that LykX subunits of all dimers were matched in 3D. Positions of the remaining Sym10 subunits were different from each other, and we utilized this difference for clustering. For each pair of dimers, we calculated two measures of dissimilarity. For two dimers, the first measure was the angle formed by two lines drawn from the center of LykX towards two centers of Sym10. The second measure was the minimal correlation between X, Y, and Z coordinates of two Sym10 subunits. For each measure, we introduced a threshold: dimers were considered similar if the angle was lower than 45, and the minimal correlation was higher than 0.5. Then, we performed clustering based on the binary similarity matrix and found groups containing at least four pairwise similar dimers. Some of these groups had a non-empty intersection; hence, we merged them until the independent clusters were obtained. As was expected, several dimers were not clustered. For each cluster, we randomly chose a typical dimer and checked whether the mutual orientation of LykX and Sym10 domains and possible orientation of dimers to the cytoplasmic membrane were biologically justified.

Typical dimers for clusters were relaxed in an orthorhombic box for 50 ns in a TIP4P solvent by use of MD, Desmond-v5.4 package (Bowers et al., 2006) of Schrödinger 2018-2. The minimal initial distance between protein and simulation box border was 20 Å. Protein dimer poses obtained after trajectory clustering were employed for ligand docking assays. To identify binding energy between LykX and Sym10 proteins, the MM-GBSA approach was applied (Schrodinger, 2013).

### Molecular dynamics parameters

MD was carried out in the Desmond-v5.4 package of Schrödinger 2018-2, using the TIP4P water solvent model; systems were neutralized by adding single-charged ions (Na^+^ or Cl^-^). In addition, 0.15 M of NaCl was added to emulate standard cytoplasmic ion concentration. MD properties were the following: ensemble class: NPT, thermostat method Nose-Hoover chain, barostat method Martyna-Tobias-Klein, relaxation time 5 ps, temperature 300 K, pressure 1,013 bar, interaction cutoff radius 9 Å. MD trajectories were clustered by the Desmond Trajectory Clustering package.

### Stability of the protein dimers in the cytoplasmic membrane model

As all three LykX-Sym10 dimers were stable in the solvent, we randomly selected one (the Afghan variant) to test its conformation stability in solvent together with the membrane. To assemble the membrane, transmembrane domains, and dimer into one complex, we performed the following steps. First, we modeled hydrophobic α-helices of LykX and Sym10 transmembrane domains and docked them to imitate the interaction between transmembrane domains in the LykX-Sym10 heterodimer. Second, we sewed subunits of LykX-Sym10 dimer to corresponding transmembrane domains via native peptide linkers through peptide bonds. Lastly, we applied MD in the solvent with the presence of the 1,2-dimyristoyl-sn-glycero-3-phosphorylcholine (DMPC) full-atom membrane model for 100 ns. The protein–lipid complex was placed in an orthorhombic box with the minimal distance between protein and simulation box borders equal to 30 Å.

### Ligand structure preparation

We considered four types of NF structures according to the possible repertoire produced by *Rhizobium leguminosarum bv. viciae*. The analysis of NF conformer structures and their stability was carried out hierarchically. First, molecular mechanics (MM) was employed to perform the initial screening of NF conformers using the MMFF94 force field (Halgren, 1996) as implemented in the Frog2 program package (Miteva et al., 2010) and the 100 lowest-energy conformers were selected for each type of NF molecule. Second, an initial assessment of each conformer’s stability in aqueous medium was carried out using the primitive implicit solvent model (COSMO). Third, based on the assessments, each NF conformer was refined by semi-empirical electronic structure calculations at the PM6-DH2X level (Řezáč and Hobza, 2011) using the MOPAC2016 program package (Stewart, 2016). For each NF, the 15 most stable conformers were then used in the docking procedure and in thermochemical analysis.

### Ligand Docking

For ligand docking, the Glide-v7.9 package of Schrödinger 2018-2 was applied (Friesner et al., 2004). We ran two flexible XP dockings for each previously obtained typical dimer; grid boxes were centered on #44 and #76 polymorphic amino acids of the LykX protein. The internal size of each grid box was 30×30×30 Å, and the external size was 50⨯50⨯50 Å. Intramolecular hydrogen bonds were rewarded, and stereochemical transitions in ligand rings were strictly forbidden. Docking results contained the ten best docking poses for each ligand. Docking poses were clusterized using the interaction fingerprints method, and the best pose for each ligand was selected by Glide Emodel criteria. Protein–ligand complexes with the best docking energy (Glide E) were chosen for each protein dimer pose.

### Relaxation and analysis of protein-ligand complexes

Sym10-LykX-NF complexes were placed in an orthorhombic box and relaxed for 300 ns using the MD approach. The minimal distance between protein and simulation box borders was 20 Å. Obtained MD trajectories were clusterized by the Desmond Trajectory Clustering method. The MM–GBSA method was applied before and after MD to identify binding energy between (i) LykX and Sym10 proteins and (ii) Sym10-LykX dimers and NF ligands.

### Quantum mechanical parameters

Quantum mechanical gas-phase calculations were carried out using the ORCA program package (ver. 4.0.1) (Neese, 2018). All structures were re-optimized using the higher-level composite RI-B97-3c method (Brandenburg et al., 2018). The energies of solvated structures were estimated by applying the hybrid COSMO-RS solvation model (Klamt and Eckert, 2004) using the CosmoTherm 16 program (Grimme, 2012) The COSMO-RS correction was applied to the DFT results obtained at the BP86/TZVP level of theory (Becke, 1988; Weigend and Ahlrichs, 2005) as a recommended procedure for the COSMO-RS calculations (Hellweg and Eckert, 2017).

## Results

### Sequence analysis

95 sequences of *P. sativum* LykX gene contained 23 polymorphic sites; the difference between two sequences was 4.7 amino acid positions on average. Among 23 sites, only four demonstrated polymorphisms between pea alleles: amino acids #44, #45, #75 and #76 (Figure 1a). Some of these four positions may potentially be responsible for the difference in European and Afghan pea phenotypes. It should be noted that European and Tajik LykX alleles are more similar to each other than to the Afghan allele variant. For modeling, we took the most frequent LykX sequence within each of the three groups of alleles.

Analysis of Sym10 sequences revealed only one valine-isoleucine polymorphism at #221 position, which was not specific for any pea allele, and valine-isoleucine substitution that was almost insignificant at both structural and chemical levels. This allowed us to conclude that the Sym10 gene sequence is universal for Afghan, Tajik, and European alleles, and guided us to use one Sym10 protein model in all further experiments.

A comparative table of polymorphisms in LykX and Sym10 amino acid sequences is presented in Supplementary Table 3.

### Protein model reconstruction

As templates for modeling, we utilized three crystal structures of three plant chitin receptors from the PDB database: 5JCD, 4EBZ, and 5LS2. The similarity values between target protein sequences (LykX or Sym10) and all templates were higher than 18% (Supplementary Table 4), but the highest values were observed for the 5LS2 template in all cases. Therefore, we used this template to model both LykX and Sym10.

We performed modeling by three different servers: the I-Tasser method (threading), the SWISS-MODEL pipeline (homology modeling), and the Phyre2 algorithm (homology modeling and threading). Analysis of Ramachandran plots and βααβ motifs of LysM domains revealed that the I-Tasser server predicted the most appropriate structures of both LykX and Sym10 receptors. After fold recognition, all models were relaxed by the energy minimization approach in Schrödinger using the implicit GB/SA water solvent model.

We constructed LykX models of European, Afghan, and Tajik alleles separately and observed that amino acids polymorphisms in the variants had no significant impact on both secondary and 3D structures of LysM domains (Figure 1c).

### LykX–Sym10 dimer assembly

We performed protein–protein docking of LykX and Sym10 proteins for each of three *P. sativum* alleles; in each case, we obtained 30 configurations of LykX-Sym10 complexes. Then, 90 models (3×30) were clustered and four distinct clusters consisting of 39 dimers in total were found (Supplementary Figure 1). After that, clusters with biologically inadequate dimer subunit orientations were filtered out. Only one cluster (Supplementary Figure 1) had dimers that met all necessary conditions. This cluster contained dimers of all three pea alleles.

From the remaining cluster, we randomly chose dimers corresponding to each pea subpopulation (IDs: A01, T00, E25 in Supplementary Data 1) and relaxed these variants in water using MD for 50 ns. Analysis of trajectories and Gibbs energy demonstrated the stability of the dimer that allowed us to consider this dimer as potentially existing in the solvent.

### Membrane model emulation

The simulation trajectory contained two clusters. For both clusters, the RMSD from the initial geometry was around 6 Å, with flexible linker fragments providing the main contribution. The closest distance between transmembrane domains was 10.8 Å, which corresponds to the semi-bounded condition for transmembrane domains (Polyansky et al., 2012). The structure of the dimer was stable during the simulation, and moved closer to the membrane after the relaxation of linker fragments (Figure 2b). Therefore, we can conclude that the obtained orientation of subunits in the LykX-Sym10 dimer may exist.

**Figure 2.**
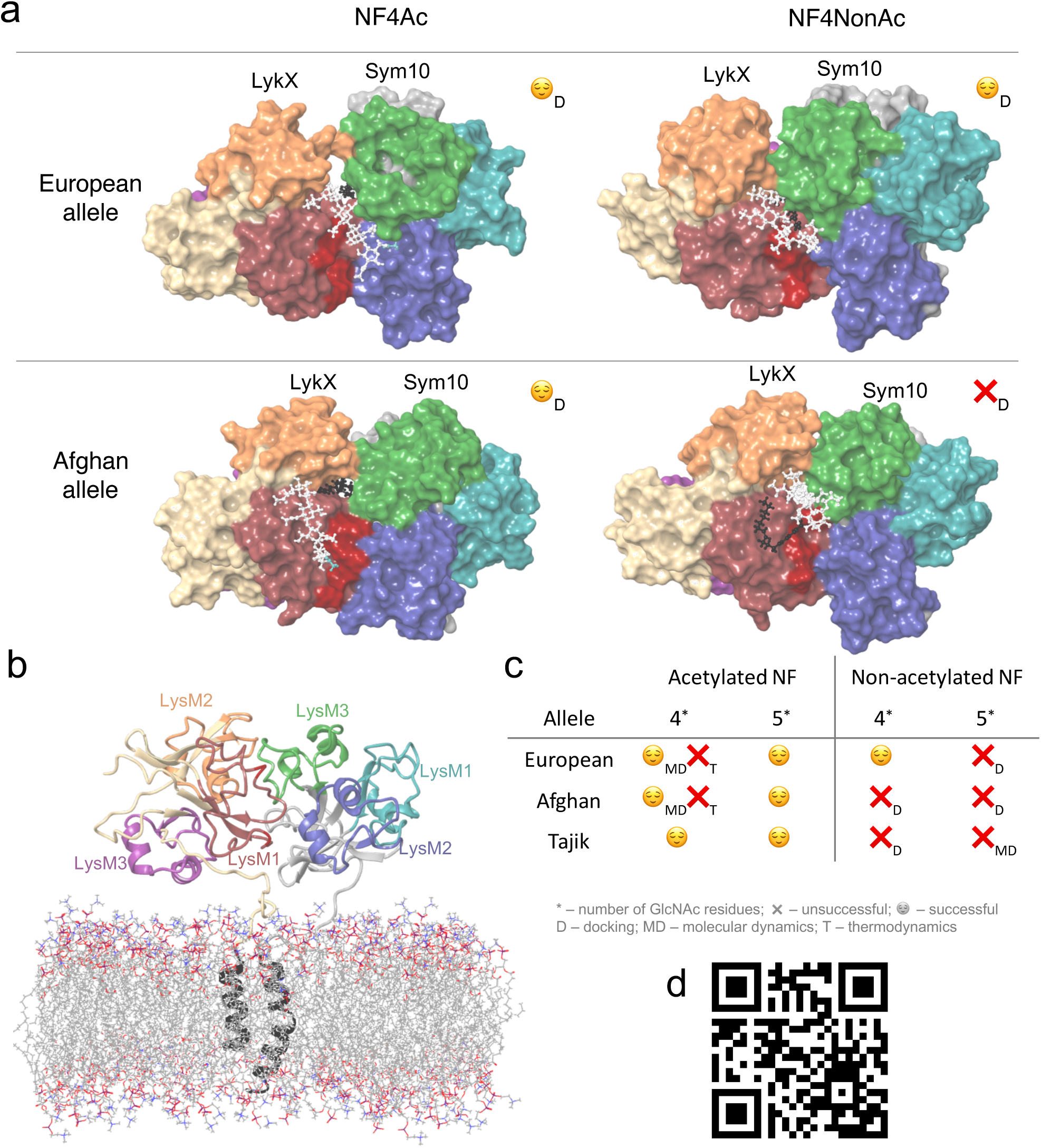
(a) Four docking poses after docking. NFs core is colors with white, fatty tail - with black, acetylation – with blue. (b) LyxX-Sym10 dimer with transmembrane domains and membrane after 100ns MD; (c) Table with successful and unfavorable NF-LyxX-Sym10 complexes; (d) QR-code to the video with the MD simulation of the NF5Ac-LykX-Sym10 complex (Tajik alleles); NPT ensemble, 300K, 500ns, TIP4P water molecules are hidden.

### Protein–ligand docking

To determine the mechanism of NF reception, all four NF types were separately docked into three dimers corresponding to three *P. sativum* alleles. We considered a dimer to recognize a NF if the fatty acid was caught in the hydrophobic pocket (cave) between the LykX and Sym10 proteins. Otherwise, plant root cells could not distinguish NFs from chitin and chitosan oligomers and trigger the early symbiotic cascades instead of an immune response. We propose that the hydrophobic recognition of the NF fatty tail by the pocket occurs at the structural level. Therefore, any structural variability in the NF tail (i.e., chain length or double bond positions) can potentially have an impact on NF–dimer specificity. This conclusion is in line with previous results, where the correspondence between population diversities of the rhizobial nodA gene (coding for the acyltransferase involved in the fatty acid tail decoration of the NF) and plant heterodimeric receptor genes was demonstrated (Igolkina et al., 2019). Plausible mechanisms of this linkage, namely coordinated variability of amino acids in contact with the surface of the receptor dimer adjacent to the NF fatty tail, were proposed (Igolkina et al., 2018). Other earlier works (Debellé et al., 1996; Moulin et al., 2004; Ritsema et al., 1996; Wu et al., 2011) also showed that nodA gene variability relates to the bacterial host range specificity. Summarizing the above, we can assume that coordinated variability of the fatty tail structure and receptor regions interacting with the fatty tail is one of the possible ways to fine-tune host specificity in legume–rhizobial symbiosis.

Based on protein–ligand docking results, NF5Ac and NF4Ac were recognized by all three dimers; NF5NonAc and NF4NonAc were recognized only by Tajik and European dimers, respectively, while Afghan dimers did not recognize any non-acetylated NFs (Figure 2a, Figure 2c). The total number of successful NF binding is 8. In each of these complexes, at least four NF–protein hydrogen bonds were formed, except T-NF5NonAc, in which only three bonds were detected (Supplementary Figure 2). LykX protein played the primary role in NF recognition, while Sym10 was involved in ligand reception only in T-NF{4,5}Ac and E-NF4NonAc complexes and seemed to act as a supporter kinase. We analyzed common amino acid residues contacting the NF and observed that Asn48 of the LykX protein was involved in NF recognition in at least five out of eight poses. At the same time, between-allele polymorphic positions in LykX — #44 and #45 — interacted with NF in the most poses. Polymorphic LykX residue #75 partly interacted with NF5Ac in Afghan and European dimers. This residue was also located in a close proximity to NFs in all other docking poses, influencing NF recognition indirectly. Some LykX amino acids were principal for the interaction with specific NF types, e.g. Asn59 took part in NF5Ac reception by both Afghan and Tajik dimers, and Thr47 was essential for NF4Ac recognition by European and Afghan dimers. The hydrophobic pocket for NF fatty tail perception was found to be formed mostly by Val37, Met38, Pro39, Ala40, Phe41, Leu42, Leu43, Tyr119, and Ala121 residues of LykX and Val217 and Phe218 residues of the Sym10 protein.

### MD relaxation of protein-ligand complexes

Along the MD trajectories, all NF-LykX-Sym10 complexes stably interacted with NF in the following manner: the NF fatty acyl tail was located in a hydrophobic pocket between dimer subunits (except T-NF5NonAc), and the NF chitin core was abundantly hydrated, forming many water bridges to dimer residues (Figure 2d).

The principal residues involved in NF reception before and after MD were similar (Supplementary Table 5). The Asn48 of the LykX protein was principal for NF recognition in all performed MD, hence we can consider it as one of the key residues in NF reception, along with #44 and #45 polymorphic residues. The role of #75 and #76 polymorphic residues in the LykX protein is less evident; they do not form contacts with NFs during MD, but could be critical in early stages of NF recognition, forming the steric landscape on the dimer surface. The Phe41 residue of LykX interacted with the fatty acyl group in all MD. The contribution of other amino acids (Tyr35, Met38, and Leu42) from the hydrophobic pocket was less pronounced and could vary.

To additionally prove the relevance of predicted NF-dimer complexes, we examined the possible mechanism of NF reception directly from the solvent. As previously demonstrated, the NF–dimer contact zone is composed of two different parts: hydrophilic contact with the NF core and hydrophobic contact with a fatty tail. However, the question remained as to what occurs first. We supposed that the primary contacts between NF and dimer are hydrophilic contacts, as they are (i) more probable in solvent and (ii) localized on the dimer surface. After the hydrophilic contacts are established, the NF fatty tail penetrates the hydrophobic pocket if their structures fit together. For this experiment, we placed NF5Ac close to the Tajik dimer surface at the previously discovered docking site nearby and MD was performed for 500 ns. We observed T-NF5Ac complex formation within 20 ns. During the first stage, the NF hydrophilic core was caught by a few separate non-stable hydrogen bonds and the fatty acid was pushed by the water molecules into the hydrophobic cave formed between both proteins. This led to the small rearrangement of LykX-Sym10 complex geometry and the effective recognition of the NF hydrophilic region by protein dimer (Supplementary Video). Thus, the principal role of LykX-Sym10 amino acid interface and the fatty acid in NF reception was proven.

### Thermochemical verification of ligand–receptor complex formation

During the formation of a ligand–receptor complex, NF changes its conformation in solvent from the free state into the binding pose. To estimate the thermodynamic benefit and principal opportunity of this transition, we computed the lowest energies of NF geometries in several states using the quantum mechanical calculations (Figure 3). The “stable” state describes the lowest energy NF conformation in gas (vacuum). The “active” NF state is obtained after docking with a dimer with further re-optimization in vacuum. For both states, we added solvent corrections, re-optimized NF conformations, and obtained “stable solvated” and “active solvated” states, respectively. We also introduced “bonded” energy, representing “active solvated” energy accounting for the docking energy (Figure 3). For each state, we calculated ΔG energy, set the “stable solvated” state as the reference point, and compared all other states with this reference point. As a result, we obtained ΔΔG energies reflecting energies of states subtracting ΔG (“stable solvated”). We then applied the following filtration criteria: negative “bonded” energy corresponds to potentially active NF molecules, while its positive value indicates impossible states. Based on these criteria, six out of eight successful docking poses were considered to be thermodynamically advantageous for forming ligand–receptor complexes. Two complexes containing acetylated NF with four chitins in the backbone — E-NF4Ac and A-NF4Ac — were filtered out (Supplementary Figure 3).

**Figure 3.**
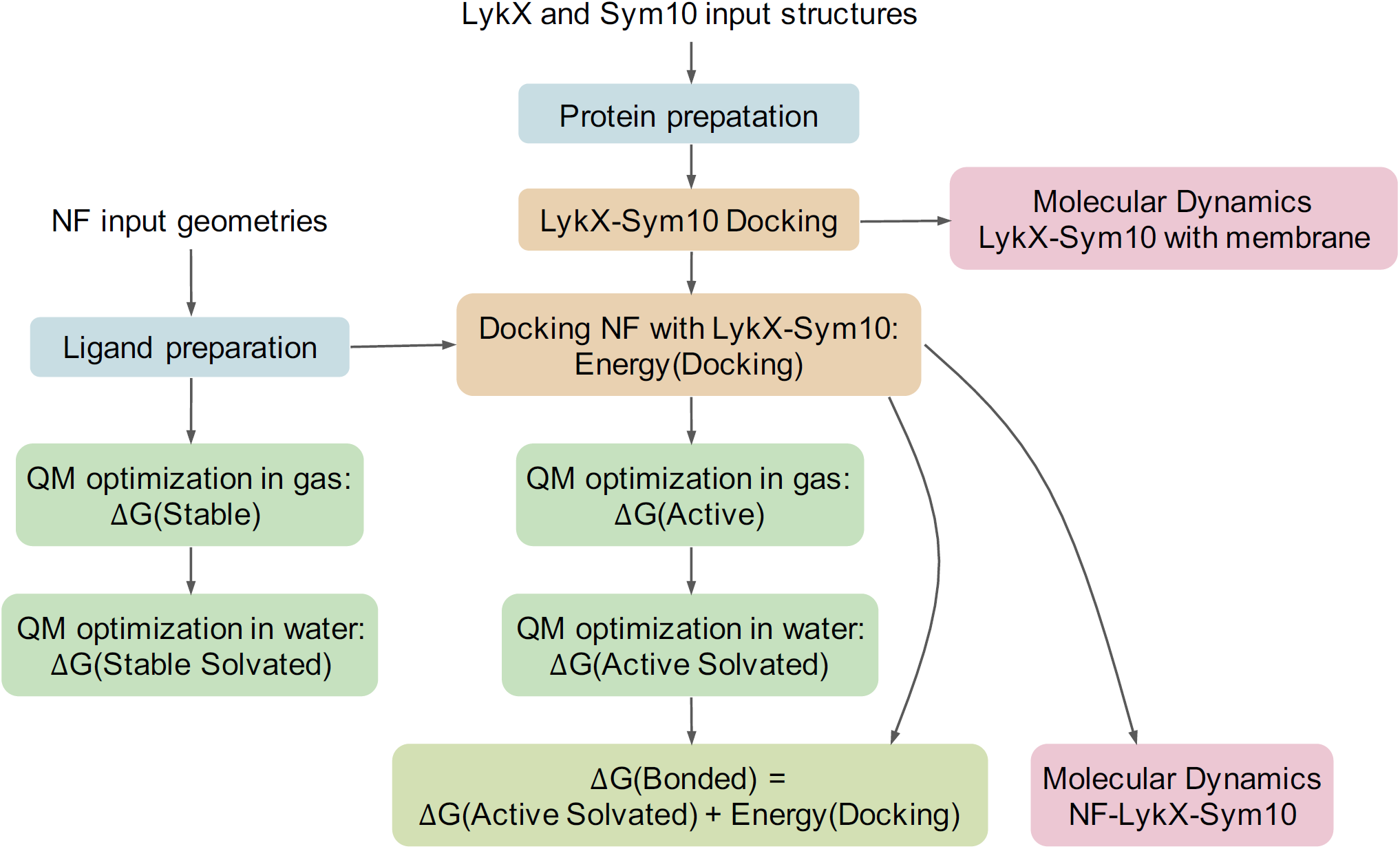
Pipeline of the analysis. Same colors highlight technically or logically similar steps.

## Discussion

We studied a possible source of variability in pea symbiotic phenotypes while interacting with *rhizobia*. In brief, (i) in addition to the European phenotype, the non-classical Afghanistan symbiotic pea phenotype was identified in 1978 (LIE, 1978); (ii) in 2003 it was first shown that legume receptors recognize rhizobial NFs working in pairs (Radutoiu et al., 2003), and (iii) after a relatively long lag, a putative pair of NF receptors in pea — LykX and Sym10 — was suggested based on sequence comparative analysis (Sulima et al., 2017).

The proposed pair of receptors requires experimental confirmation, for example, by studying mutants of LykX (currently in progress, Zhukov, V., personal communication). However, this approach may only justify the involvement of the target gene in controlling the Afghanistan phenotype, but may not validate the direct interaction between the mutant LykX-Sym10 receptor and NFs. Experimental detection of the NF-LykX-Sym10 interaction is difficult because of challenges in isolating the receptor heterodimer in its natural form.

Due to the complexity of the system under study, molecular modeling is a promising way to test whether the LykX-Sym10 heterodimer is the NF receptor responsible for European/Afghanistan symbiotic phenotypes. We considered four stages in the assembly of the NF-LykX-Sym10 complex and performed molecular modeling of each of them (Figure 3):

1. stability of the LykX-Sym10 heterodimer in solvent in the presence of a membrane by MD;
2. successful docking of NF to LykX-Sym10;
3. stability of the NF-LykX-Sym10 complex in solvent during MD;
4. possibility of NF-LykX-Sym10 complex formation in terms of thermodynamic benefits.

We not only demonstrated that LykX-Sym10 is stable in solvent in the presence of a membrane, but also found that LykX-Sym10 heterodimers of both European and Afghanistan phenotypes have the same 3D configuration. The natural and non-deleterious amino acid polymorphisms in receptors should not significantly impact their mutual disposition in a correctly working dimer. Therefore, the observed match in dimer structures likely indicates that the obtained LykX-Sym10 configuration is correct. Moreover, this configuration corresponds to the previously proposed stable “sandwich-like” configuration (Igolkina et al., 2018), where a LysM domain of one subunit binds between LysM2 and LysM3 domains of another subunit. This 3D conservatism of the LykX-Sym10 structure could be a criterion for filtering legume receptor dimers in other modeling studies.

Analysis of NF docking poses in LykX-Sym10 revealed that in most cases (8 out of 12), the NF fatty acid tail was located in the hydrophobic pocket formed by the contact zone of two receptor subunits. This result is also in line with the previously elucidated stable configuration (Igolkina et al., 2018). Therefore, we considered these eight poses to be biologically justified. It should be noted that only non-acetylated NFs were involved in four unsuccessful docking poses.

For each of eight NF-LykX-Sym10 complexes we applied both MD and thermodynamic analysis, each playing the filtration role. It is important to emphasize that these steps are independent and answer different questions: (i) whether the complex is stable, and (ii) whether the complex is possible, respectively. Sets of NF-LykX-Sym10 complexes, which were filtered out by each procedure, were not overlapped, which demonstrates that MD and thermodynamic calculations cannot be interchangeable steps.

During the thermodynamic calculations, we compared ΔG energies of NF in different states assuming that biologically active NF geometries correspond to stable states. The chemical structure of NF makes it very flexible, which leads to the plurality of conformations. We assumed that NF stable states could be biologically active if their free energies allow them to interact with the dimer. This assumption is in line with our observation about the conservatism of LykX-Sym10 configurations. Summarizing all together, we can speculate that the conservatism or stability of molecular structures is the important property for biological functioning, and should be taken into account during filtration of biologically justifying models.

Only five out of twelve NF-LykX-Sym10 complexes successfully passed all steps of the analysis (Figure 2c), and they precisely match the known pea symbiotic differences: peas of European phenotype can form a symbiosis with *Rhizobium leguminosarum bv. viciae*, producing both acetylated and non-acetylated NFs, while peas of Afghanistan phenotype react on only acetylated NFs. It is important to note that all the LykX-Sym10 heterodimer complexes considered here formed successful complexes with NF5Ac. Based on our analysis and match between estimated and observed pea phenotypes, we can conclude that the LykX gene is a suitable candidate gene for Sym2.

In sum, molecular modeling has shown itself to be an effective way of functional analysis complementary to traditional approaches, and its results provide appealing directions for further studies in this area, such as determining functional meaning of minor components of NF mixtures produced by *rhizobia*, or reconstruction of the evolutionary history of signal systems in plant– microbe interactions.

## Supporting information

LykX_Sym10_protein_dimers, protein homology modeling geometries

## Acknowledgements

This work has been supported by the Russian Science Foundation # 19-16-00081 (conceptualisation, introduction and conclusion) and # 17-76-30016 (Design of allele set for the molecular modelling, PsSym10 alleles sequencing). Quantum chemical calculations were carried out using SurfSara supercomputer facilities with a support from NWO.

## Contributions

All authors read and approve of the final manuscript. Solovev Ya.V. and Igolkina A.A. contributed equally. The methodology was developed by E.E.A., Ya.V.S., A.A.I., and Yu.B.P.; data analysis was performed by Ya.V.S., A.A.I., and P.O.K.; visualization was performed by Ya.V.S. and A.A.I.; 3D modelling and QM calculations were performed by Ya.V.S. and P.O.K. Initial writing and draft preparation was done by Ya.V.S. and A.A.I.; review and editing were made by all authors; data is curated by A.S.S. and V.A.Z; software is provided by Yu.B.P. and E.A.P.

## Supplementary material

**Supplementary Figure 1.**
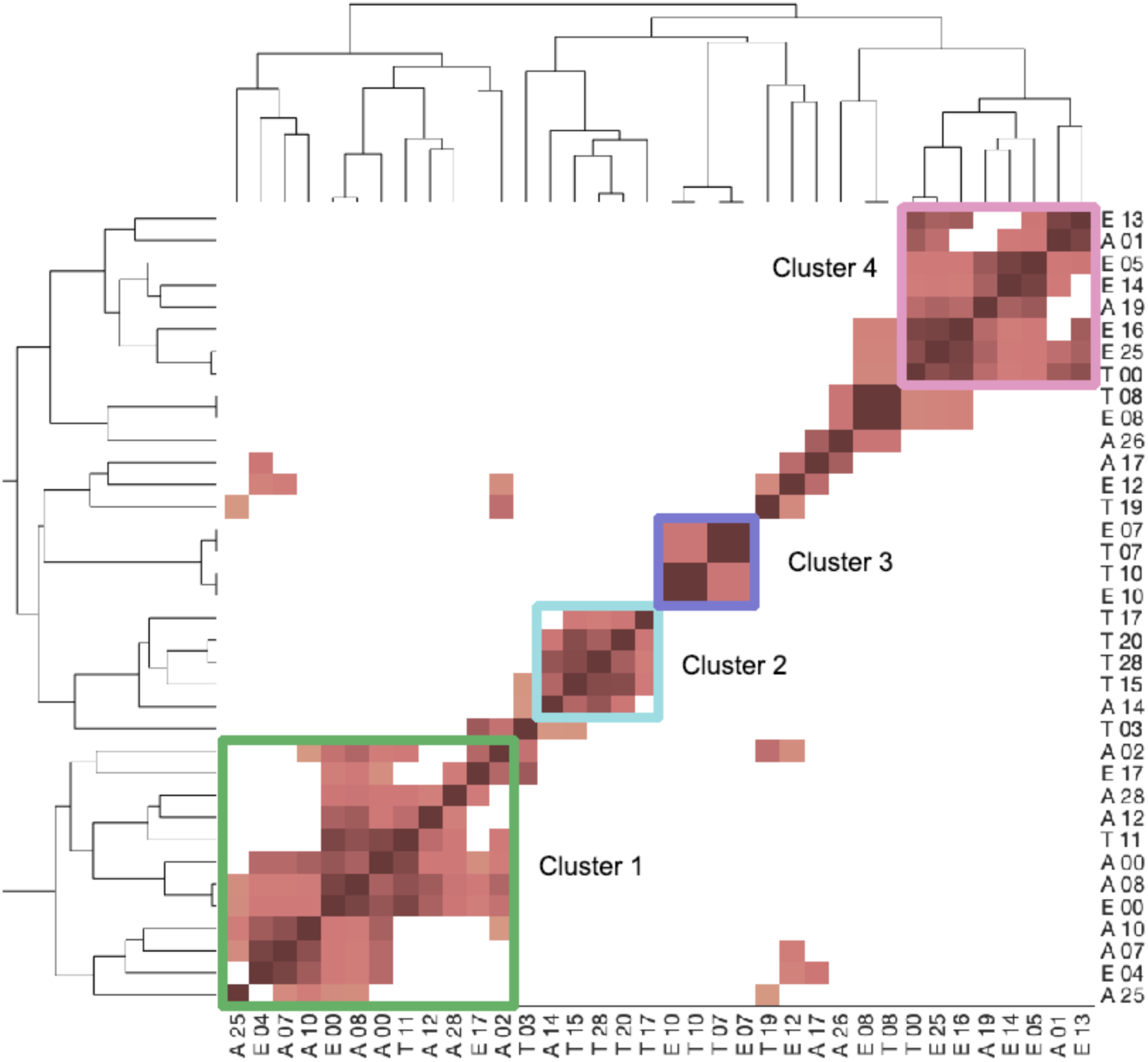
Similarity of mutual orientation between LykX and Sym10 receptors in dimers. Four separate clusters were identified and highlighted with colors. Afghan alleles are denoted with A, Tajik – with T, and European – with E.

**Supplementary Figure 2.**
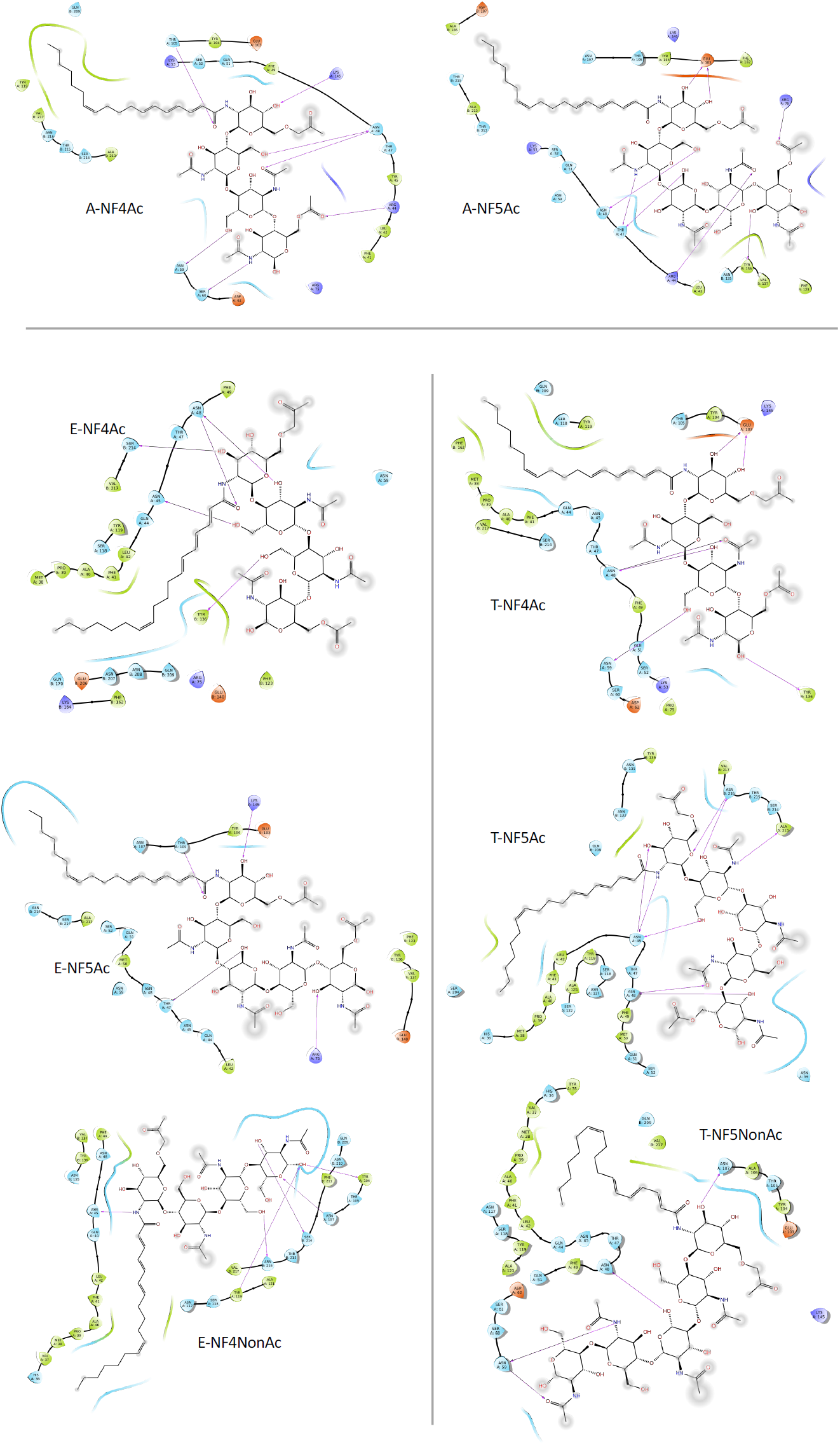
Protein-ligand interactions in 8 LyxX-Sym10-NFs complexes. Hydrogen bonds are marked as purple arrow lines.

**Supplementary Figure 3.**
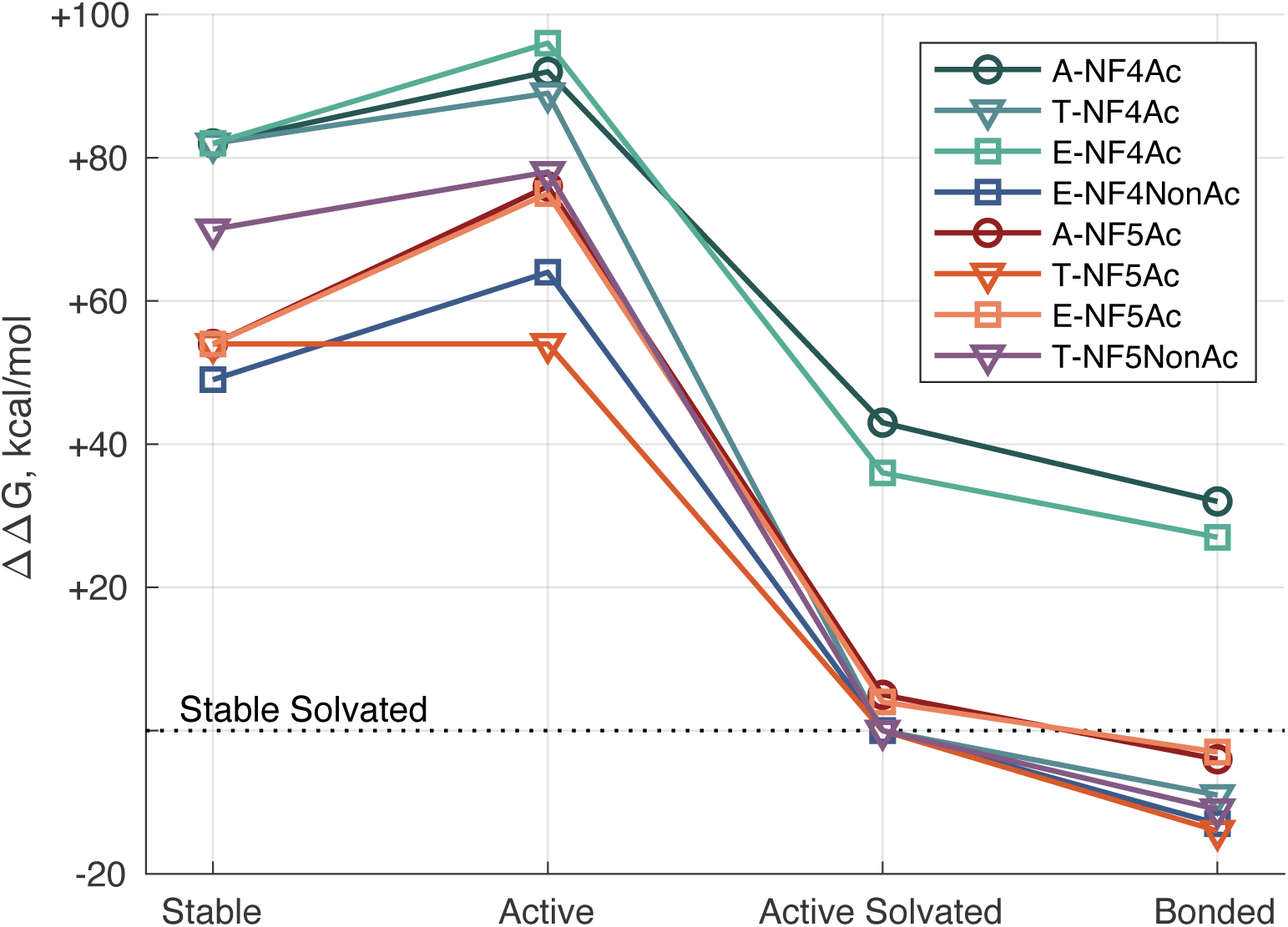
*ΔΔ*G relative free energy plot for LykX-Sym10-NFs complexes in different computational conditions. The 0-value at Y-axis corresponds to the Stable Solvated energy as a reference level. Energies in other conditions are presented as deviations from the reference.

**Supplementary Table 1.** *P. sativum* receptor genes annotation

| Receptor gene | Allele | Accession numbers | References | Amount |
| --- | --- | --- | --- | --- |
| *PsSym10* |  |  |  | 8 |
|  | EU | AJ575250 | Madsen et al., 2003 |  |
|  | EU | AJ575251 | Madsen et al., 2003 |  |
|  | EU | AJ575252 | Madsen et al., 2003 |  |
|  | EU | AJ575253 | Madsen et al., 2003 |  |
|  | EU | MN727811 | This work |  |
|  | AF | MN727808 | This work |  |
|  | AF | MN727809 | This work |  |
|  | AF | MN727810 | This work |  |
| *PsLykX* |  |  |  | 95 |
|  | EU | MF155382-MF155468 | Sulima et al., 2017 | 87 |
|  | AF | MN187362 | Sulima et al., 2019 |  |
|  | TJ | MN187363 | Sulima et al., 2019 |  |
|  | AF | MN200353 | Sulima et al., 2019 |  |
|  | AF | MN200354 | Sulima et al., 2019 |  |
|  | AF | MN200355 | Sulima et al., 2019 |  |
|  | TJ | MN200356 | Sulima et al., 2019 |  |
|  | AF | MN200357 | Sulima et al., 2019 |  |
|  | AF | MN200358 | Sulima et al., 2019 |  |

**Supplementary Table 2.**
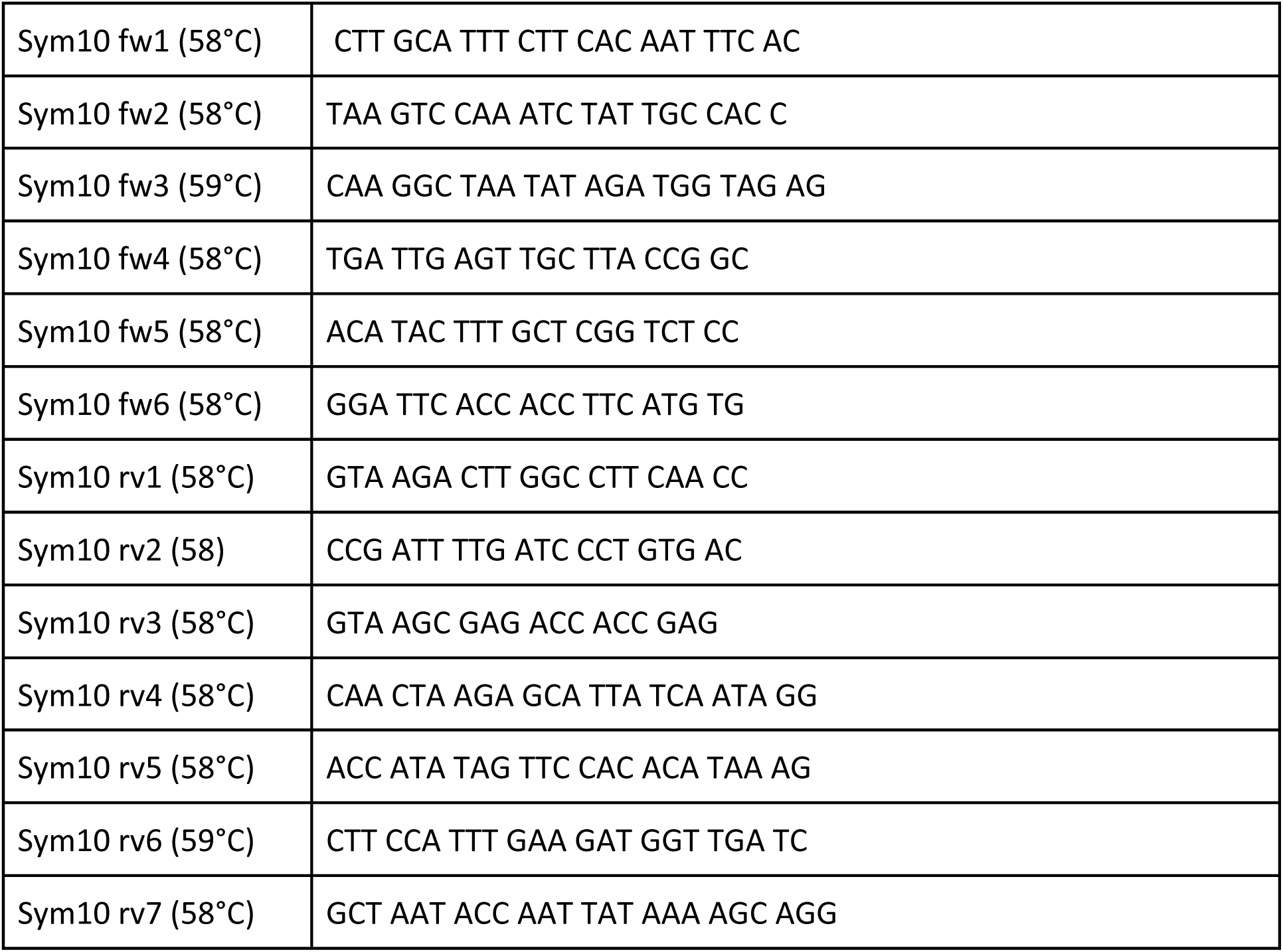
Primers used for sequencing of PsSym10 alleles (5’-3’) and corresponding temperature.

**Supplementary Table 3.** Amino acid polymorphism in the LykX protein of the European, Tajik, and Afghan subpopulations. In positions #44, #45, #75, and #76 subpopulation-specific differences are marked by bold, other variable amino acids in polymorphic sites are separated by ‘∼’ symbol. In position #45 both asparagine (N) and serine (S) are relevant for European subpopulation, but serine was presented only in the single sequence.

| Index | Number of position | Domain | Variable amino acids |  |  | Blosum 90 matrix |
| --- | --- | --- | --- | --- | --- | --- |
|  |  |  | Afghanistan | Tajik | European |  |
| 1 | 5 | Signal peptide | F | F | F ~ L | 0 |
| 2 | 9 | Signal peptide | L | L | F ~ L | 0 |
| 3 | 13 | Signal peptide | L | L | V ~ L | 4 |
| 4 | 14 | Signal peptide | E | E ~ D | E | 3 |
| 5 | 16 | Signal peptide | V | V | V ~ F | -1 |
| 6 | 18 | Signal peptide | F | F | S ~ F | -3 |
| 7 | 23 | Unstructured region | K | K | K ~ Q | 1 |
| 8 | 42 | LysM1 | K | K | K ~ L | -3 |
| 9 | 44 | LysM1 | R | Q | Q | 1 |
| 10 | 45 | LysM1 | Y | N | N ~ S | -5 / -3 |
| 11 | 48 | LysM1 | N | N | N ~ K | 0 |
| 12 | 75 | LysM1 | R | P | R | -2 |
| 13 | 76 | LysM1 | D | A | A | -3 |
| 14 | 82 | LysM1 | F | F | S ~ F | -3 |
| 15 | 86 | LysM1 | I | I | V ~ I | 5 |
| 16 | 111 | LysM2 | T | T | T ~ S | 3 |
| 17 | 128 | LysM2 | V | V ~ F | V | -1 |
| 18 | 134 | LysM2 | S | S | G ~ S | 1 |
| 19 | 136 | LysM2 | H | H | H ~ D | -4 |
| 20 | 142 | LysM2 | I | A ~ I | A ~ I | -2 |
| 21 | 183 | LysM3 | A | A | A ~ I | -2 |
| 22 | 184 | LysM3 | S | S | S ~ F | -3 |
| 23 | 191 | LysM3 | I | I | M ~ I | 3 |

**Supplementary Table 4.** Protein amino acid sequences similarity in percent (%) between three model proteins and four *P. sativum* model proteins. The 5LS2 and the 4EBZ sequences are significantly closer to each other and to target proteins than the 5JCD sequence. The highest sequence similarity with target proteins was detected for the 5LS2 model, its values are marked by bold.

| Model | 4EBZ | 5JCD | 5LS2 |
| --- | --- | --- | --- |
| Afghanistan_Sym10 | 23.6% | 18.7% | **23.9%** |
| European_Sym10 | 22.9% | 18.3% | **24.4%** |
| Afghanistan_LykX | 45.2% | 18.0% | **49.5%** |
| European_LykX | 44.7% | 18.6% | **48.9%** |

**Supplementary Table 5.** The most crucial amino acids involved in dimer-NF interactions during MD simulations. Data is presented as percentage of trajectory duration.

| Amino acid | A-NF4Ac | A-NF5Ac | E-NF4Ac | E-NF4NonAc | E-NF5Ac | T-NF4Ac | T-NF5Ac | T-NF5NonAc |
| --- | --- | --- | --- | --- | --- | --- | --- | --- |
| LykX_103 | 94% | 14% | 0% | 2% | 70% | 86% | 0% | 61% |
| LykX_104 | 13% | 27% | 34% | 22% | 24% | 23% | 8% | 24% |
| LykX_105 | 80% | 52% | 23% | 34% | 99% | 95% | 8% | 97% |
| LykX_107 | 1% | 38% | 38% | 44% | 0% | 0% | 10% | 61% |
| LykX_119 | 17% | 30% | 20% | 11% | 25% | 15% | 24% | 59% |
| LykX_40 | 2% | 2% | 62% | 9% | 2% | 2% | 68% | 0% |
| LykX_41 | 29% | 22% | 33% | 40% | 24% | 31% | 14% | 45% |
| LykX_42 | 0% | 0% | 22% | 31% | 23% | 1% | 31% | 0% |
| LykX_44 | 39% | 8% | 24% | 6% | 42% | 83% | 19% | 8% |
| LykX_45 | 19% | 56% | 88% | 67% | 100% | 75% | 75% | 7% |
| LykX_47 | 65% | 8% | 4% | 1% | 77% | 64% | 0% | 71% |
| LykX_48 | 94% | 77% | 81% | 55% | 72% | 78% | 20% | 81% |
| LykX_49 | 31% | 32% | 2% | 46% | 9% | 28% | 4% | 22% |
| LykX_52 | 77% | 2% | 0% | 0% | 50% | 86% | 0% | 55% |
| LykX_59 | 45% | 1% | 1% | 0% | 12% | 61% | 1% | 71% |
| Sym10_132 | 0% | 16% | 0% | 40% | 62% | 0% | 14% | 0% |
| Sym10_135 | 0% | 0% | 11% | 68% | 41% | 8% | 6% | 0% |
| Sym10_136 | 2% | 1% | 65% | 49% | 74% | 17% | 56% | 0% |
| Sym10_137 | 0% | 0% | 53% | 16% | 35% | 12% | 0% | 0% |
| Sym10_211 | 18% | 0% | 24% | 43% | 0% | 2% | 4% | 46% |
| Sym10_213 | 9% | 0% | 29% | 68% | 4% | 0% | 43% | 3% |
| Sym10_214 | 0% | 0% | 82% | 71% | 52% | 0% | 24% | 2% |
| Sym10_215 | 0% | 32% | 6% | 46% | 6% | 0% | 26% | 0% |
| Sym10_216 | 0% | 74% | 57% | 61% | 81% | 0% | 64% | 0% |
| Sym10_217 | 2% | 18% | 21% | 8% | 17% | 2% | 39% | 2% |

## References

Arrighi J-F. 2006. The Medicago truncatula Lysine Motif-Receptor-Like Kinase Gene Family Includes NFP and New Nodule-Expressed Genes. PLANT Physiol. doi: 10.1104/pp.106.084657

Baldwin IL, Fred EB, Hastings EG. 1927. Grouping of Legumes According to Biological Reactions of Their Seed Proteins. Possible Explanation of Phenomenon of Cross Inoculation. Bot Gaz. doi: 10.1086/333728

Becke AD. 1988. Density-functional exchange-energy approximation with correct asymptotic behavior. Phys Rev A. doi: 10.1103/PhysRevA.38.3098

Bensmihen S, de Billy F, Gough C. 2011. Contribution of NFP LysM domains to the recognition of Nod factors during the Medicago truncatula/Sinorhizobium meliloti symbiosis. PLoS One. doi: 10.1371/journal.pone.0026114

Bowers KJ, Chow E, Xu H, Dror RO, Eastwood MP, Gregersen BA, Klepeis JL, Kolossvary I, Moraes MA, Sacerdoti FD, Salmon JK, Shan Y, Shaw DE. 2006. Scalable algorithms for molecular dynamics simulations on commodity clusters Proceedings of the 2006 ACM/IEEE Conference on Supercomputing, SC’06. doi: 10.1145/1188455.1188544

Bozsoki Z, Cheng J, Feng F, Gysel K, Vinther M, Andersen KR, Oldroyd G, Blaise M, Radutoiu S, Stougaard J. 2017. Receptor-mediated chitin perception in legume roots is functionally separable from Nod factor perception. Proc Natl Acad Sci. doi: 10.1073/pnas.1706795114

Brandenburg JG, Bannwarth C, Hansen A, Grimme S. 2018. B97-3c: A revised low-cost variant of the B97-D density functional method. J Chem Phys. doi: 10.1063/1.5012601

Broghammer A, Krusell L, Blaise M, Sauer J, Sullivan JT, Maolanon N, Vinther M, Lorentzen A, Madsen EB, Jensen KJ, Roepstorff P, Thirup S, Ronson CW, Thygesen MB, Stougaard J. 2012. Legume receptors perceive the rhizobial lipochitin oligosaccharide signal molecules by direct binding. Proc Natl Acad Sci. doi: 10.1073/pnas.1205171109

Buendia L, Girardin A, Wang T, Cottret L, Lefebvre B. 2018. LysM Receptor-Like Kinase and LysM Receptor-Like Protein Families: An Update on Phylogeny and Functional Characterization. Front Plant Sci. doi: 10.3389/fpls.2018.01531

Davis EO, Evans IJ, Johnston AWB. 1988. Identification of nodX, a gene that allows Rhizobium leguminosarum biovar viciae strain TOM to nodulate Afghanistan peas. MGG Mol Gen Genet. doi: 10.1007/BF00330860

Debellé F, Plazanet C, Roche P, Pujol C, Savagnac A, Rosenberg C, Promé JC, Dénarié J. 1996. The NodA proteins of Rhizobium meliloti and Rhizobium tropici specify the N-acylation of Nod factors by different fatty acids. Mol Microbiol. doi: 10.1046/j.1365-2958.1996.00069.x

Denarie J. 1996. Rhizobium Lipo-Chitooligosaccharide Nodulation Factors: Signaling Molecules Mediating Recognition and Morphogenesis. Annu Rev Biochem. doi: 10.1146/annurev.biochem.65.1.503

Firmin JL, Wilson KE, Carlson RW, Davies AE, Downie JA. 1993. Resistance to nodulation of cv. Afghanistan peas is overcome by nodX, which mediates an O-acetylation of the Rhizobium leguminosarum lipo-oligosaccharide nodulation factor. Mol Microbiol. doi: 10.1111/j.1365-2958.1993.tb01961.x

Friesner RA, Banks JL, Murphy RB, Halgren TA, Klicic JJ, Mainz DT, Repasky MP, Knoll EH, Shelley M, Perry JK, Shaw DE, Francis P, Shenkin PS. 2004. Glide: A New Approach for Rapid, Accurate Docking and Scoring. 1. Method and Assessment of Docking Accuracy. J Med Chem. doi: 10.1021/jm0306430

Grimme S. 2012. Supramolecular binding thermodynamics by dispersion-corrected density functional theory. Chem - A Eur J. doi: 10.1002/chem.201200497

Halgren TA. 1996. Merck molecular force field. I. Basis, form, scope, parameterization, and performance of MMFF94. J Comput Chem. doi: 10.1002/(SICI)1096-987X(199604)17:5/6<490::AID-JCC1>3.0.CO;2-P

Hellweg A, Eckert F. 2017. Brick by brick computation of the gibbs free energy of reaction in solution using quantum chemistry and COSMO-RS. AIChE J. doi: 10.1002/aic.15716

Igolkina AA, Bazykin GA, Chizhevskaya EP, Provorov NA, Andronov EE. 2019. Matching population diversity of rhizobial nodA and legume NFR5 genes in plant–microbe symbiosis. Ecol Evol. doi: 10.1002/ece3.5556

Igolkina AA, Porozov YB, Chizhevskaya EP, Andronov EE. 2018. Structural Insight Into the Role of Mutual Polymorphism and Conservatism in the Contact Zone of the NFR5–K1 Heterodimer With the Nod Factor. Front Plant Sci. doi: 10.3389/fpls.2018.00344

Kelley LA, Mezulis S, Yates CM, Wass MN, Sternberg MJE. 2015. The Phyre2 web portal for protein modeling, prediction and analysis. Nat Protoc. doi: 10.1038/nprot.2015.053

Kirienko AN, Porozov YB, Malkov N V., Akhtemova GA, Le Signor C, Thompson R, Saffray C, Dalmais M, Bendahmane A, Tikhonovich IA, Dolgikh EA. 2018. Role of a receptor-like kinase K1 in pea Rhizobium symbiosis development. Planta. doi: 10.1007/s00425-018-2944-4

Kirienko AN, Vishnevskaya NA, Kitaeva AB, Shtark OY, Kozyulina PY, Thompson R, Dalmais M, Bendahmane A, Tikhonovich IA, Dolgikh EA. 2019. Structural variations in LysM domains of LysM-RLK psK1 may result in a different effect on Pea–Rhizobial symbiosis development. Int J Mol Sci. doi: 10.3390/ijms20071624

Klamt A, Eckert F. 2004. Prediction of vapor liquid equilibria using COSMOtherm. Fluid Phase Equilib. doi: 10.1016/j.fluid.2003.08.018

Kozakov D, Brenke R, Comeau SR, Vajda S. 2006. PIPER: An FFT-based protein docking program with pairwise potentials. Proteins Struct Funct Genet. doi: 10.1002/prot.21117

Kozik A, Heidstra R, Horvath B, Kulikova O, Tikhonovich I, Ellis THN, van Kammen A, Lie TA, Bisseling T. 1995. Pea lines carrying syml or sym2 can be nodulated by Rhizobium strains containing nodX; sym1 and sym2 are allelic. Plant Sci. doi: 10.1016/0168-9452(95)04123-C

Kozik A, Matvienko M, Scheres B, Paruvangada VG, Bisseling T, Van Kammen A, Ellis THN, LaRue T, Weeden N. 1996. The pea early nodulin gene PsENOD7 maps in the region of linkage group I containing sym2 and leghaemoglobin. Plant Mol Biol. doi: 10.1007/BF00020614

Kumar S, Stecher G, Li M, Knyaz C, Tamura K. 2018. MEGA X: Molecular evolutionary genetics analysis across computing platforms. Mol Biol Evol. doi: 10.1093/molbev/msy096

Lie TA. 1978. Symbiotic specialisation in pea plants: The requirement of specific Rhizobium strains for peas from Afghanistan Annals of Applied Biology. doi: 10.1111/j.1744-7348.1978.tb00743.x

Liu S, Wang J, Han Z, Gong X, Zhang H, Chai J. 2016. Molecular Mechanism for Fungal Cell Wall Recognition by Rice Chitin Receptor OsCEBiP. Structure. doi: 10.1016/j.str.2016.04.014

Liu T, Liu Z, Song C, Hu Y, Han Z, She J, Fan G, Wang J, Jin C, Chang J, Zhou JM, Chai J. 2012. Chitin-induced dimerization activates a plant immune receptor. Science (80-). doi: 10.1126/science.1218867

Madhavi Sastry G, Adzhigirey M, Day T, Annabhimoju R, Sherman W. 2013. Protein and ligand preparation: Parameters, protocols, and influence on virtual screening enrichments. J Comput Aided Mol Des. doi: 10.1007/s10822-013-9644-8

Mergaert P, Van Montagu M, Holsters M. 1997. Molecular mechanisms of Nod factor diversity. Mol Microbiol. doi: 10.1111/j.1365-2958.1997.mmi526.x

Miteva MA, Guyon F, Tufféry P. 2010. Frog2: Efficient 3D conformation ensemble generator for small compounds. Nucleic Acids Res. doi: 10.1093/nar/gkq325

Moulin L, Béna G, Boivin-Masson C, Stȩpkowski T. 2004. Phylogenetic analyses of symbiotic nodulation genes support vertical and lateral gene co-transfer within the Bradyrhizobium genus. Mol Phylogenet Evol. doi: 10.1016/S1055-7903(03)00255-0

Neese F. 2018. Software update: the ORCA program system, version 4.0. Wiley Interdiscip Rev Comput Mol Sci. doi: 10.1002/wcms.1327

Ovtsyna AO, Rademaker G-J, Esser E, Weinman J, Rolfe BG, Tikhonovich IA, Lugtenberg BJJ, Thomas-Oates JE, Spaink HP. 2007. Comparison of Characteristics of the nodX Genes from Various Rhizobium leguminosarum Strains. Mol Plant-Microbe Interact. doi: 10.1094/mpmi.1999.12.3.252

Ovtsyna AO, Rademaker GJ, Esser E, Weinman J, Rolfe BG, Tikhonovich IA, Lugtenberg BJJ, Thomas-Oates JE, Spaink HP. 1999. Comparison of characteristics of the nodX genes from various Rhizobium leguminosarum strains. Mol Plant-Microbe Interact. doi: 10.1094/MPMI.1999.12.3.252

Polyansky AA, Volynsky PE, Efremov RG. 2012. Multistate organization of transmembrane helical protein dimers governed by the host membrane. J Am Chem Soc. doi: 10.1021/ja303483k

Provorov NA. 1994. The interdependence between taxonomy of legumes and specificity of their interaction with rhizobia in relation to evolution of the symbiosis. Symbiosis.

Radutoiu S, Madsen LH, Madsen EB, Felle HH, Umehara Y, Grønlund M, Sato S, Nakamura Y, Tabata S, Sandal N, Stougaard J. 2003. Plant recognition of symbiotic bacteria requires two LysM receptor-like kinases. Nature 425:585–592. doi: 10.1038/nature02039

Radutoiu S, Madsen LH, Madsen EB, Jurkiewicz A, Fukai E, Quistgaard EMH, Albrektsen AS, James EK, Thirup S, Stougaard J. 2007. LysM domains mediate lipochitin-oligosaccharide recognition and Nfr genes extend the symbiotic host range. EMBO J. doi: 10.1038/sj.emboj.7601826

Řezáč J, Hobza P. 2011. A halogen-bonding correction for the semiempirical PM6 method. Chem Phys Lett. doi: 10.1016/j.cplett.2011.03.009

Ritsema T, Wijfjes AHM, Lugtenberg BJJ, Spaink HP. 1996. Rhizobium nodulation protein NodA is a host-specific determinant of the transfer of fatty acids in Nod factor biosynthesis. Mol Gen Genet. doi: 10.1007/BF02174343

Rogers SO, Bendich AJ. 1985. Extraction of DNA from milligram amounts of fresh, herbarium and mummified plant tissues. Plant Mol Biol. doi: 10.1007/BF00020088

Roos K, Wu C, Damm W, Reboul M, Stevenson JM, Lu C, Dahlgren MK, Mondal S, Chen W, Wang L, Abel R, Friesner RA, Harder ED. 2019. OPLS3e: Extending Force Field Coverage for Drug-Like Small Molecules. J Chem Theory Comput. doi: 10.1021/acs.jctc.8b01026

Rudemo M. 2000. Statistical Shape Analysis. I. L. Dryden and K. V. Mardia, Wiley, Chichester 1998. No. of pages: xvii+347. Price: £60.00.ISBN 0-471-95816-6. Stat Med. doi: 10.1002/1097-0258(20001015)19:19<2716::aid-sim590>3.0.co;2-o

Schrödinger. 2018. Protein Preparation Wizard | Schrödinger. Schrödinger Release 2018-1.

Schrodinger LLC. 2013. MacroModel, Version 10.2. New York.

Sears OH, Carroll WR. 1927. Cross inoculation with cowpea and soybean nodule bacteria. Soil Sci. doi: 10.1097/00010694-192712000-00003

Stewart JJP. 2016. MOPAC2016. Stewart Comput Chem. doi: 10.2106/JBJS.G.00147

Sulima AS, Zhukov VA, Afonin AA, Zhernakov AI, Tikhonovich IA, Lutova LA. 2017. Selection Signatures in the First Exon of Paralogous Receptor Kinase Genes from the Sym2 Region of the Pisum sativum L. Genome. Front Plant Sci. doi: 10.3389/fpls.2017.01957

Sulima AS, Zhukov VA, Kulaeva OA, Vasileva EN, Borisov AY, Tikhonovich IA. 2019. New sources of Sym2A allele in the pea (Pisum sativum L.) carry the unique variant of candidate LysM-RLK gene LykX. PeerJ. doi: 10.7717/peerj.8070

Viprey V, Rosenthal A, Broughton WJ, Perret X. 2000. Genetic snapshots of the Rhizobium species NGR234 genome. Genome Biol. doi: 10.1186/gb-2000-1-6-research0014

Waterhouse A, Bertoni M, Bienert S, Studer G, Tauriello G, Gumienny R, Heer FT, De Beer TAP, Rempfer C, Bordoli L, Lepore R, Schwede T. 2018. SWISS-MODEL: Homology modelling of protein structures and complexes. Nucleic Acids Res. doi: 10.1093/nar/gky427

Weigend F, Ahlrichs R. 2005. Balanced basis sets of split valence, triple zeta valence and quadruple zeta valence quality for H to Rn: Design and assessment of accuracy. Phys Chem Chem Phys. doi: 10.1039/b508541a

Wu LJ, Wang HQ, Wang ET, Chen WX, Tian CF. 2011. Genetic diversity of nodulating and non-nodulating rhizobia associated with wild soybean (Glycine soja Sieb. & Zucc.) in different ecoregions of China. FEMS Microbiol Ecol. doi: 10.1111/j.1574-6941.2011.01064.x

Xiao TT, Schilderink S, Moling S, Deinum EE, Kondorosi E, Franssen H, Kulikova O, Niebel A, Bisseling T. 2014. Fate map of Medicago truncatula root nodules. Dev. doi: 10.1242/dev.110775

Zhang Y. 2009. I-TASSER: Fully automated protein structure prediction in CASP8. Proteins Struct Funct Bioinforma. doi: 10.1002/prot.22588

Zipfel C, Oldroyd GED. 2017. Plant signalling in symbiosis and immunity. Nature. doi: 10.1038/nature22009

